# EPI distortion correction for simultaneous human brain stimulation and imaging at 3T

**DOI:** 10.1101/547935

**Authors:** Hyuntaek Oh, Jung Hwan Kim, Jeffrey M. Yau

**Author notes:** Corresponding author: Jeffrey M. Yau, One Baylor Plaza, T111, Houston, TX 77030.

## Abstract

Transcranial magnetic stimulation (TMS) can be paired with functional magnetic resonance imaging (fMRI) in simultaneous TMS-fMRI experiments. These multimodal experiments enable causal probing of network architecture in the human brain which can complement alternative network mapping approaches. Critically, merely introducing the TMS coil into the scanner environment can sometimes produce substantial magnetic field inhomogeneities and spatial distortions which limit the utility of simultaneous TMS-fMRI. We assessed the efficacy of point spread function corrected echo planar imaging (PSF-EPI) in correcting for the field inhomogeneities associated with a TMS coil at 3T. In phantom and brain scans, we quantitatively compared the coil-induced distortion artifacts measured in PSF-EPI scans to artifacts measured in conventional echo-planar imaging (EPI) and a simultaneous multi-slice sequence (SMS)-EPI. While we observed substantial coil-related artifacts in the data produced by the conventional EPI and SMS sequences, PSF-EPI produced data that had significantly greater signal-to-noise and less distortions. In phantom scans with the PSF-EPI sequence, we also characterized the temporal profile of dynamic artifacts associated with TMS delivery and found that image quality remained high as long as the TMS pulse preceded the RF excitation pulses by at least 50ms. Lastly, we validated the PSF-EPI sequence in human brain scans involving TMS and motor behavior as well as resting state fMRI scans. Our collective results demonstrate the superiority of PSF-EPI over conventional EPI and SMS sequences for simultaneous TMS-fMRI when coil-related artifacts are a concern. The ability to collect high quality resting state fMRI data in the same session as the simultaneous TMS-fMRI experiment offers a unique opportunity to interrogate network architecture in the human brain.

## Introduction

Transcranial magnetic stimulation (TMS) is a non-invasive approach for manipulating human brain activity (Hallet, 2007). Because TMS-induced activity in a targeted region can lead to activity changes in remote but connected regions, combining TMS with functional magnetic resonance imaging (fMRI) or electroencephalographic (EEG) enables the characterization of functional and distributed cortical networks through the causal manipulation of human brain activity (Shastri et al., 1999; Baudewig et al., 2000; Bestmann et al., 2004; Ruff et al., 2006, 2007; Hanakawa et al., 2009; Blankenburg et al., 2010; Peters et al., 2012; Bestmann & Feredoes, 2013; Leitão et al., 2015, 2017). A major advantage of simultaneous TMS-fMRI over other network mapping approaches like resting state fMRI and diffusion tensor imaging is the ability to characterize state-dependent changes in network architecture (Bestmann et al., 2005; Blankenburg et al., 2008; Feredoes et al., 2011; Leitão et al., 2013, 2015; Rahnev et al., 2016; Ruff et al., 2006; Sack et al., 2006, 2007).

Historically, a major challenge with simultaneous TMS-fMRI is field map inhomogeneity, generated by the introduction of the TMS coil into the MRI environment, which produces spatial distortions and signal loss in the acquired images. Importantly, the presence and magnitude of coil-related artifacts may differ dramatically depending on the particular hardware used in the experiments. When coil-related artifacts are present, the spatial distortions in the images depend on the orientation of image slices with respect to the orientation of the TMS coil (Baudewig et al., 2000; Bestmann et al., 2003a): Substantially greater geometric distortions and ghost artifacts are present when the frequency-encoding gradient is perpendicular to the TMS coil plane as compared to when the frequency-encoding direction and slice orientation are parallel to the TMS coil. Accordingly, the parallel alignment scheme has been used as a common strategy to reduce coil-related artifacts in simultaneous TMS-fMRI experiments (Bestmann et al., 2003a; Moisa et al., 2009; Weiskopf et al., 2009; Bungert et al., 2012; Navarro de Lara et al., 2015, 2017). However, this strategy has limitations. First, there is no automated and reliable method for optimally aligning functional prescriptions with the TMS coil even with the use of vitamin capsules (Bestmann et al., 2006; Sack et al., 2007) or water-filled tubes (Bestmann et al., 2003a; Leitã o et al., 2012; Weijer et al., 2014) to visualize the TMS coil. Moreover, though using a parallel alignment can improve signal quality over the brain generally, this strategy may actually cause a greater signal reduction in specific brain regions. For instance, a consequence of aligning the slice prescriptions to the TMS coil plane can be an exacerbation of the signal loss that is already prominent in areas such as the orbitofrontal cortices (Weiskopf et al., 2006; Moisa et al., 2009). Lastly, susceptibility artifacts may persist even after achieving parallel alignment between the frequency-encoding direction and the TMS coil.

When using coils, stimulators, and filtering systems that produce artifacts, alternative hardware-based strategies have also been employed to treat the static artifacts and signal loss associated with the introduction of the TMS coil to the scanner environment. A relay-diode combination can be effective in reducing image artifacts caused by leakage currents (Weiskopf et al., 2009). Susceptibility artifacts can also be reduced through passive shimming (Bungert et al., 2012). Furthermore, customized radiofrequency coil arrays may be used to improve MR sensitivity (Navarro de Lara et al., 2015, 2017). These strategies require specialized hardware, though, and they can sometimes increase ghosting artifacts while also attenuating TMS intensities.

While much of the efforts to improve data quality in simultaneous TMS-fMRI experiments have focused on procedural strategies or hardware innovations, there have been limited efforts to address TMS-induced static field artifacts using specialized pulse sequences, though such sequences have been developed as general solutions to correct for susceptibility artifacts. Point spread function (PSF) corrected echo planar imaging (PSF-EPI) has been used to minimize susceptibility artifacts and distortion, which is a critical problem particularly with ultra-high field imaging (Robson et al., 1997; Zeng & Constable., 2002; Zaitsev et al., 2004; Chung et al., 2011; In & Speck., 2012; Oh et al., 2012; In et al., 2015, 2016, 2017a; Loureiro et al., 2017). In brief, PSF mapping is based on the combination of EPI phase-encoding gradients (which are vulnerable to distortions) with spin-warp phase-encoding gradients (which are geometrically accurate). PSF mapping provides a summary of the EPI distortions – caused by B_0_ field inhomogeneity, susceptibility, chemical shift, or eddy currents – which can then be corrected. In addition to its use in correcting for image stretching and compression at ultra-high fields, PSF-EPI has also been used in fMRI scans with deep brain stimulation devices to minimize the metal-induced distortion near the implants (In et al., 2017b).

Here, we tested the utility of PSF-EPI for simultaneous TMS-fMRI performed at 3T. In phantom and brain scans, we measured and compared the signal distortion and loss associated with the mere introduction of a TMS coil into the scanner environment in images acquired using PSF-EPI, conventional EPI, and multi-band simultaneous multi-slice (SMS)-EPI (Setsompop et al., 2012). In addition to characterizing the static artifacts related to the presence of the TMS coil, we also characterized the acute artifacts associated with delivery of TMS pulses before and during image acquisition. We validated the PSF-EPI sequence in an in vivo experiment by quantifying fMRI BOLD signal changes associated with TMS application and the performance of a motor task. Lastly, we acquired resting state fMRI datasets using the PSF-EPI sequence and characterized functional connectivity patterns with the TMS coil positioned over different regions. These collective experiments enabled us to establish the utility of the PSF-EPI sequence for simultaneous TMS-fMRI experiments in which coil-related artifacts impact image quality. Our finding that high quality resting state fMRI data can be acquired along with simultaneous TMS-fMRI data using the same experimental setup establishes the feasibility of relating these complementary approaches for characterizing network architecture in the human brain.

## Materials and methods

### General overview

We performed 5 separate experiments. Experiment 1 was aimed at characterizing the static spatial artifacts in phantom scans acquired using PSF-EPI and conventional EPI generated by the introduction of a TMS coil into the scanner in the absence TMS coil discharging. Experiment 2 was aimed at characterizing the same static artifacts in human brain scans acquired using PSF-EPI, conventional EPI, and SMS-EPI sequences. Experiment 3 was aimed at characterizing the acute artifacts in images acquired with PSF-EPI associated with delivery of a single TMS pulse at different intensities and times relative to image acquisition at multiple coil orientations. Experiment 4 was aimed at validating PSF-EPI for simultaneous TMS-fMRI by characterizing BOLD signal changes associated with the performance of a cued motor task and TMS application over sensorimotor cortex. Experiment 5 was aimed at characterizing resting state functional connectivity (RSFC) using the PSF-EPI sequence in the presence of the TMS coil to test whether consistent connectivity patterns could be attained with the coil positioned over different brain regions.

### Magnetic resonance imaging

Functional and structural imaging data were acquired on a Siemens MAGNETOM Trio 3T scanner (Siemens Medical Solutions, Erlangen, Germany) using the posterior elements (6 channels) of the 12-channel head matrix coil (part #08622644) and the flexible the 6-channel body matrix coil (part #08622651). Note that this particular combination and configuration of coil arrays may be differentially vulnerable to TMS-related artifacts compared to alternative rigid or flexible head coils and depending on the specific TMS system used. This issue is beyond the scope of our current study, as we focused on characterizing the relative benefits of PSF-EPI over conventional EPI and SMS-EPI sequences for a particular setup; however, the topic of imaging coil designs for simultaneous TMS-fMRI has been extensively considered previously (Navarro De Lara et al., 2015). In all experiments, we first performed a localizer scan (1.1×1×7 mm, TR/TE = 8.6/4 ms, field of view (FOV) = 250mm, flip angle = 20°, 1 slice, readout bandwidth = 320 Hz/pixel). In each session, we also always acquired functional data using the PSF-EPI sequence (3-mm isotropic voxels, TR/TE = 3000/30 ms, interleaved slice acquisition, phase oversampling = 0%, FOV = 210 mm, flip angle = 84°, 46 slices, readout bandwidth = 1374 Hz/pixel, PAT mode: GRAPPA, parallel imaging acceleration factor = 2). In Experiments 1 and 2, functional data were also acquired using a conventional EPI sequence (3-mm isotropic voxels, TR/TE = 3000/30 ms, interleaved slice acquisition, phase oversampling = 0%, FOV = 210 mm, flip angle = 84°, 46 slices, readout bandwidth = 1374 Hz/pixel, PAT mode: GRAPPA, parallel imaging acceleration factor = 2). Importantly, in both the PSF-EPI and conventional EPI sequences, all of the slices were acquired in the first 2750 ms of the 3000-ms repetition time allowing for a 250-ms gap between consecutive volumes. We additionally acquired two separate 3D gradient echo (GRE) images (2.4-mm isotropic voxels, TR = 488 ms, TE = 4.92 and 7.38 ms, FOV = 230 mm, flip angle = 40°, 45 slices, readout bandwidth = 301 Hz/pixel) and used these data to generate field maps using AFNI (Cox, 1996; Cox and Jesmanowicz, 1999) and FSL (Jenkinson et al., 2012). The field maps (Fig. 1) reveal clear susceptibility distortion in the B_0_ field in the vicinity of the TMS coil. In Experiment 2, we also acquired functional data using a SMS-EPI sequence (2 x blipped-controlled aliasing in parallel imaging (CAIPI) multi-slice acceleration (shift factor 4) with GRAPPA (acceleration factor 2) (Breuer et al., 2005; Setsompop et al., 2012); 3-mm isotropic voxel resolution, TR/TE = 1480/30 ms, phase oversampling = 0%, FOV = 210 mm, flip angle = 70°, 46 slices, readout bandwidth = 1488 Hz/pixel). To enable direct comparisons between the functional datasets, the prescriptions of the conventional EPI and SMS-EPI sequences were set to be identical to or closely matching those of the PSF-EPI sequence, respectively. We acquired T1-weighted structural images using a 3D FLASH sequence (1.6 x 1.6 x 3 mm^3^, TR/TE = 6.62/1.85 ms, FOV = 210 mm, flip angle = 15°, 48 slices, readout bandwidth = 130 Hz/pixel, parallel imaging acceleration factor = 2). Matching the prescriptions of the 3D FLASH scans to the functional scans enabled a direct comparison between the EPI scans that are vulnerable to field inhomogeneities to a T1-weighted sequence that is not. In sessions involving brain scans, we also acquired structural images using a magnetization prepared rapid gradient echo (MPRAGE) sequence (1 mm isotropic voxel resolution, TR/TE = 2600/3.02 ms, FOV = 256 mm, flip angle = 8°, 176 slices, readout bandwidth = 130 Hz/pixel, parallel imaging acceleration factor = 2). These structural data were used for estimating the TMS target site, data visualization, co-registration with the functional data from the simultaneous TMS-fMRI and resting state fMRI experiments.

**Fig. 1.**
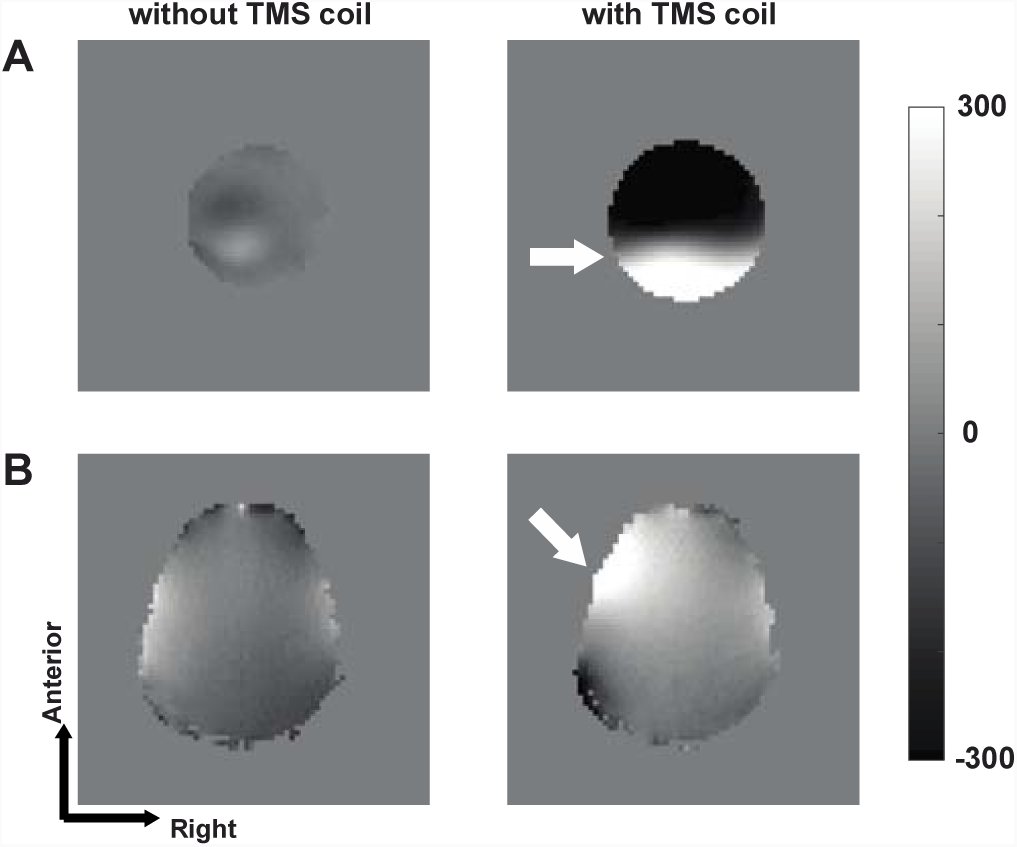
(A) Field map of a spherical phantom without and with the TMS coil. (B) Field map of brain without and with the TMS coil. The field maps were calculated in units of rad/s. The TMS coil (white arrow) was positioned superior to the phantom with its plane aligned to the transverse imaging plane and over the left sensorimotor cortex.

### Transcranial magnetic stimulation

A MRI-compatible figure-of-eight coil (Magstim Co. UK; inner diameter: 42mm; outer diameter: 80mm; number of turns: 10; cable length: 5.75 m to the filter box) was connected to 2 Magstim Rapid^2^ stimulator units located outside the MRI suite via a 2-to-1 filter adaptor (Magstim Co. UK). A filter box and ferrite sleeves served to attenuate radio frequency (RF) noise. The filter box (Magstim Co. UK) contains a shielded inductive filter to help prevent spurious RF interference from entering the MRI room via the stimulating coil cable. Such RF interference can come from the Magstim unit or from the antenna effect of the long cable. In addition the filter box contains a relay that can be used to short any direct current coming from the Magstim equipment itself. A custom-built platform and coil holder secured the TMS coil to the scanner bed. This platform also supported the body matrix coil that provided anterior coverage of the phantom and brain.

The TMS coil was labeled by four microcentrifuge tubes (0.5 ml) filled with 1 part Gadolinium (Gd; 1:300) and 1 part 0.9% sodium chloride solution. These Gd-filled tubes were highly visible markers that defined the position and orientation of the TMS coil in the localizer scans and in the T1-weighted scans. In Experiment 4, we localized the TMS coil and its position and orientation with respect to the brain in the T1-weighted images using a pipeline, analogous to our previously described method (Yau et al., 2013), implemented in AFNI and custom Matlab scripts (R2017a. The MathWorks, Natick, MA, USA). This procedure enables the identification of the cortical region falling immediately beneath the center of the TMS coil (Yau et al., 2013).

TMS timing was controlled by Psychtoolbox-3 (Kleiner et al., 2007) in MATLAB (2011b, MathWorks) running on a MacBook Pro (model A1278; OS X 10.9.5, 2.5 GHz Core i5, 4 GB of RAM). TMS was triggered by TTL pulses sent via a DAQ device (model USB-1208FS, Measurement Computing Corporation).

### Experiment 1. Static artifact characterization in phantom scans

In phantom scans (agarose gel 2% phantom; 48-mm circumference) acquired using PSF-EPI and conventional EPI, we quantified signal loss and image distortion artifacts induced by the TMS coil in the absence of TMS discharging. Each scan comprised 100 volumes. We scanned the phantom using each pulse sequence with and without the TMS coil. The TMS coil was positioned superior to the phantom with its plane aligned to the transverse imaging plane. To establish the dependence of the static artifacts on the relative orientation of the image slices and the TMS coil, in separate scans we prescribed functional slices that were either parallel or oblique to the TMS coil surface as guided by the Gd-filled markers (Fig. 2A). Functional data were acquired in transverse planes with the phase encoding gradients in a right-left direction. For the parallel prescription, the orientation of the image slices was aligned parallel to the TMS coil surface. For the oblique prescription, the slices were tilted approximately 15° from the parallel prescription in both the sagittal and coronal planes.

**Fig. 2.**
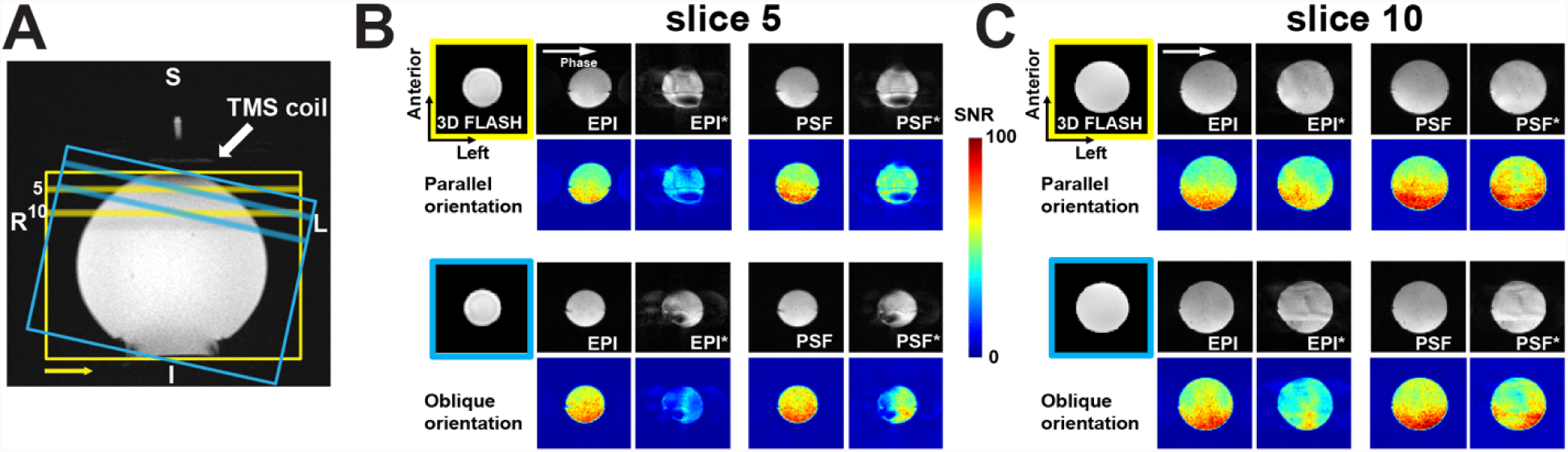
(A) Phantom measurement setup. Conventional EPI and PSF-EPI data acquired with either parallel (yellow) or oblique (blue) prescriptions with respect to the TMS coil surface. Slice prescriptions are guided by Gadolinium-filled markers labeling the TMS coil which are visible in a set of three-plane localizer scans. The phase encoding direction was a right-left direction (marked with the yellow arrow). The 5^th^ and 10^th^ slices in the image stacks are indicated by the bars. (B) Comparisons of the raw conventional EPI and PSF-EPI images and SNR maps calculated for the 5^th^ slice. Data acquired in the parallel orientation (top 2 rows) and oblique orientation (bottom 2 rows) reveal the dependence of image quality on acquisition orientation. Structural images acquired using the 3D FLASH sequence are undistorted by the TMS coil. The asterisks indicate data acquired in the presence of the TMS coil. (C) Comparisons of the 10^th^ slice. Conventions as in *B*. R = right, L = left, S = superior, I = inferior, Phase = phase-encoding direction.

### Experiment 2. Static artifact characterization in brain scans

In brain scans acquired using PSF-EPI, conventional EPI, and SMS-EPI, we quantified signal loss and image distortion artifacts induced by the TMS coil in the absence of TMS discharging. Images were only acquired with the TMS coil positioned over left sensorimotor cortex (Talairach:-51, −5, 42) (Fig. 4A). In this experiment, we did not manipulate the slice prescriptions with respect to the TMS coil: Slice acquisition orientation was maintained in the transverse plane. Each scan comprised 100 volumes.

### Experiment 3. Characterization of acute artifacts associated with TMS discharge

Eddy currents produced by TMS discharge interfere with subsequent image acquisition by causing signal loss and distortion in specific spatial-temporal patterns (Shastri et al., 1999). A common strategy for avoiding these acute artifacts is to introduce a temporal gap between TMS discharge and subsequent RF-excitation and image acquisition (Bestmann et al., 2003b; Ruff et al., 2006). In Experiment 3, we wished to characterize the acute artifact induced by a single TMS pulse in functional data acquired using PSF-EPI to establish the temporal delay required after the TMS pulse to avoid image perturbation. Critically, although the PSF-EPI sequence we used had a nominal repetition time of 3000 ms, a single volume was acquired in 2750 ms thereby leaving a 250-ms gap between consecutive volume acquisitions during which TMS could be delivered. In separate scans on a phantom, we manipulated the time of a single TMS discharge during and after this 250-ms interval and quantified its impact on image acquisition. TMS was delivered −125, −100, −75, −50, 0, and +50 ms with respect to RF excitation and image acquisition. To characterize how acute discharge-related artifacts scaled with TMS intensity, we repeated the experiment at 2 TMS intensities, expressed as a percentage of maximum stimulator output (70% and 100%). Thus, we performed 12 scans in this experiment. Each scan began with an initial 4 TMS-free volumes. Subsequently, a TMS-containing “cycle” began: A volume was acquired without TMS, a single TMS pulse was delivered at the designated time, and 3 additional volumes were acquired. This cycle repeated 10 times over the scan yielding a total of 44 volumes per TMS condition. We tested each TMS condition with the coil positioned superior or lateral to the phantom (i.e., with the TMS coil pointing in a direction aligned with or perpendicular to the B_0_ field, respectively). The slices were acquired parallel to the plane of the TMS coil in each scan.

### Experiment 4. Simultaneous TMS-fMRI with PSF-EPI

We performed a set of simultaneous TMS-fMRI scans to validate the use of the PSF-EPI sequence for measuring BOLD signal changes related to acute TMS and the performance of a motor behavior. A right-handed male (20 years old) participated in the experiment after giving the written informed consent. All procedures were approved by the Institutional Review Board for human subject research at the Baylor College of Medicine. The participant reported no history of neurological disorders or impaired sensorimotor function. Head movement was limited by foam padding and the subject wore earplugs and noise-attenuating earmuffs to minimize noise from the scanner and the click sounds associated with TMS discharge. The TMS coil was positioned over the left sensorimotor cortex (x = −51, y = −5, z = 42), and slice acquisition orientation was maintained in the transverse plane irrespective of the TMS coil position.

The primary goals of this experiment were to verify that robust BOLD signal changes could be measured in the vicinity of the TMS coil and in regions removed from the coil rather than to characterize activation patterns for the purpose of inferring network architecture. We used a block design paradigm to characterize BOLD signal changes associated with TMS and bimanual finger tapping (Fig. 8A). In separate scans, blocks consisted of either low intensity TMS (TMS_Low_; 40% maximum stimulator output), high intensity TMS (TMS_High_; 50% maximum stimulator output), or a motor task performed concurrently with low intensity TMS (FT+TMS_Low_) (Bestmann et al., 2005; Yau et al,. 2013). Note that these TMS intensities were both below the threshold required to produce visible muscle activation at rest, but subthreshold TMS can result in BOLD signal changes distributed over cortical and subcortical sensorimotor networks (Bestmann et al., 2005; Yau et al,. 2013). The motor task involved bimanual finger-tapping: The participant touched his thumbs to each finger in sequence continuously over the duration of the block (9 sec). Each block comprised 3 volumes and blocks were separated by 7 volumes during which the participant rested without receiving TMS. During each block, the subject received 3 bursts of TMS with each burst (3 pulses, 16Hz, inter-pulse interval: 62.5ms) triggered during the 250-ms gaps separating volume acquisitions. Each burst began at the onset of the inter-volume gap thereby providing 125 ms for the eddy currents to dissipate before acquisition of the subsequent volume. Each scan comprised 10 blocks (104 volumes; total scan time: 5:2 min). This scan began with 4 TMS-free volumes.

### Experiment 5. Resting state fMRI with PSF-EPI

We performed a series of resting state fMRI scans using the PSF-EPI sequence in the presence of the TMS coil. The goal of this experiment was to verify that clear resting state connectivity patterns were attainable in the presence of the TMS coil with the PSF-EPI sequence and that these patterns were invariant to the positioning of the TMS coil. In each of 3 sessions (inter-session interval: 6 days), we acquired fifteen continuous minutes of resting state fMRI data (300 volumes) in the one participant who also participated in Experiment 4. The TMS coil was positioned over different scalp locations in each session. In session 1, the TMS coil was over the left middle frontal gyrus (x = −44, y = 3, z = 54). In session 2, the coil was positioned over the right superior frontal gyrus (x = 39, y = 26, z = 49). In session 3, the coil was positioned over the left inferior parietal lobule (x = −49, y = −32, z = 50). Slice acquisition orientation was maintained in the transverse plane irrespective of TMS coil position. The subject underwent each resting state fMRI experiment with eyes closed. No TMS pulses were triggered during these scans.

### Data analysis

All data analyses were performed using AFNI and Matlab (R2018a).

### Static artifacts in phantom data

We computed 2 metrics to compare the impact of TMS coil presence and slice prescription on the PSF-EPI and conventional EPI phantom data. As a measure of signal strength, we generated temporal SNR (tSNR) maps for each sequence under each condition. For each voxel, we computed:

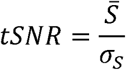

where 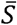 is the mean signal of the fMRI time series and σ_*s*_ is the standard deviation of the signal. To focus our analysis on the voxels contained within the phantom, we defined an analysis mask using the T1-weighted volume acquired with the 3D FLASH sequence: The mask comprised all voxels that exceeded an intensity threshold defined as 0.1 of the maximum signal in the T1-weighted volume. Because the prescriptions of the PSF-EPI and conventional EPI sequences were matched, the mask was applied identically to both datasets as we compared the sequences slice-wise. To compare the tSNR values computed for each sequence, we conducted a 3-way repeated-measures ANOVA with EPI sequence, TMS coil presence, and prescription orientation as factors. We performed post-hoc comparisons, with Bonferroni-corrections, based on significant main and interaction effects.

Because the presence of the TMS coil causes spatial distortions as well as signal drop, we compared the T1-weighted 3D FLASH volume, which is not susceptible to the presence of the TMS coil, to the mean volumes acquired with PSF-EPI and conventional EPI to characterize the relative image distortions related to the TMS coil presence and slice prescriptions in the EPI data. Using the 3D FLASH volume as a reference, we computed structure similarity (SSIM) indices for each EPI sequence (Wang et al., 2004). SSIM is a reference metric that quantifies the quality of a comparison image with respect to a reference image as a weighted combination of comparisons between the images’ luminance, contrast, and structure. Because SSIM depends on luminance, we first normalized the reference (R) and comparison (C) volumes by each volume’s maximum intensity. For each slice in the EPI volumes, we then computed:

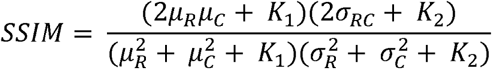

where µ_R_ is the average normalized intensity in the reference slice, µ_C_ is the average normalized intensity in the comparison slice, 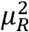 is the variance of the reference slice, 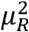 is the variance of the comparison slice, σRC is the covariance of the reference and comparison slices, and *K*_*1*_ and *K*_*2*_ are stabilizing parameters fixed at 0.01 and 0.03, respectively. SSIM values range from 0 to 1 with larger values of SSIM indicating greater similarity between the reference and comparison data. To compare the SSIM values computed for each sequence, we conducted a 3-way repeated-measures ANOVA with EPI sequence, TMS coil presence, and prescription orientation as factors. We performed post-hoc comparisons, with Bonferroni-corrections, based on significant main and interaction effects.

### Static artifacts in brain data

To compare the impact of the TMS coil on resting brain data acquired using PSF-EPI, conventional EPI, and SMS-EPI, we calculated and compared tSNR and SSIM using analogous analyses as those described for the phantom data. Note that we matched the number of volumes acquired using each sequence rather than total scan time, so the SMS-EPI scan was substantially shorter in duration than the other scans. Given this design choice, the relative tSNR calculated for the SMS-EPI data is an underestimate which could be improved with more images if the PSF-EPI, conventional EPI, and SMS-EPI sequences had been equated according to scan duration instead. The tSNR analyses only comprised voxels contained within the skull as defined in the 3D FLASH data (threshold: >10% of the maximum intensity). We visualized tSNR brain maps using previously described methods (Taylor et al., 2018). In the SSIM analyses, we first performed skull-stripping on only the 3D FLASH data (using the 3dSkullStrip function in AFNI) and then compared this reference volume to the EPI volumes without thresholding. Removing the skull from the 3D FLASH data better enabled us to account for spatial distortions in the EPI data that pushed the signals from the brain outside of the skull. Because we only measured signals in the presence of the TMS coil and with a single prescription, we performed one-way repeated measures ANOVA with EPI sequence as the factor.

### Acute artifacts in phantom data

To quantify acute artifacts in the PSF-EPI sequence related to TMS discharge, we compared volumes acquired during or subsequent to TMS delivery to a reference volume acquired before any TMS discharges. We used the 4^th^ volume in the scan as the reference. Because magnetic fields produced by TMS decay over space and time, we computed the root mean square error (RMSE) between corresponding slices in the reference and comparison volumes for each TMS pulse time, as done previously (Navarro de Lara et al., 2017). Note that we excluded data from a single slice measurement from the analysis because its signal intensity differed from the mean of the other 9 slices acquired under identical conditions by −33 times the standard deviation across the 9 slices. To test if TMS coinciding with RF-excitation results in altered signal intensities in subsequent acquisitions, as shown previously (Bestmann et al., 2003a), we also assessed potential signal changes in the 2^nd^ and 3^rd^ volumes acquired after TMS delivery.

### Simultaneous TMS-fMRI data

Data analysis for the simultaneous TMS-fMRI experiment was performed in volume space using AFNI. As we have done previously (Pérez-Bellido et al., 2017), we performed data preprocessing including slice timing correction, motion correction, despiking, and coregistration with the structural data. The functional data were spatially smoothed by convolution with a 4-mm FWHM isotropic Gaussian kernel after spatial normalization into the Talairach space. We analyzed the data by fitting a general linear model that included 3 regressors of interest corresponding to the TMS_Low_, TMS_High_, and FT+TMS_Low_ blocks convolved with gamma-variate functions. Head motion parameters and drift parameters (linear, quadratic, cubic, and quartic) were included as nuisance regressors. To localize regions whose signals changed in association with finger-tapping and low-intensity TMS, statistical results for the FT+TMS_Low_>baseline contrast were thresholded at a false discovery rate (FDR) corrected q < 0.05 over the whole brain. To localize regions within the sensorimotor system (established from the FT+ TMS_Low_ responses) whose signals changed in association with TMS, we contrasted TMS_Low_ and TMS_High_ against baseline (FDR q < 0.05, masked by FT+TMS_Low_).

### Resting state fMRI data

Data analysis for the resting state fMRI experiment was performed in surface space using AFNI. A surface model of the brain was created using FreeSurfer (Dale et al., 1999). We performed standard data preprocessing (slice timing correction, motion correction, and despiking) before projecting the data into surface space. A bandpass filter (0.01 −0.1 Hz) was applied to reduce the effect of low and high frequency physiological noise (Bright & Murphy, 2015). The data were spatially smoothed by convolution with a 4-mm FWHM isotropic Gaussian kernel in surface space. For each session separately, the data in each hemisphere were sorted into 180 discrete parcels (Glasser et al., 2016). We calculated the mean time series in each parcel and computed the correlation between the mean time series in each pair of parcels to generate a RSFC matrix for each session.

As a first step in determining the overall similarity of the RSFC patterns across sessions, we calculated the pairwise correlations between the upper triangle of the RSFC matrices generated for the sessions (Gordon et al., 2017). To determine whether the TMS coil influenced RSFC patterns specifically in regions near the coil, we computed the RSFC profile 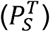 for the parcel identified as each session’s TMS target site (*T*) and compared these RSFC profiles across sessions (*S*). For instance, we identified the parcel corresponding to the session 1 TMS target (i.e., the left middle frontal gyrus) and computed the RSFC strength between this parcel and all parcels across the two hemispheres for in the session 1 data to define a vector of 360 correlation values, 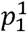 We then calculated the RSFC profiles for the same parcel using the session 2 and session 3 data to define 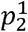 and 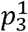 If the TMS coil distorted the resting state data locally near the coil, we might expect that the RSFC profiles for the target parcel *T* would differ across sessions in which the coil was moved. To quantify RSFC profile similarity for the same parcel across sessions, we calculated the pairwise correlations between 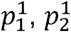, and 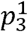 and averaged the correlation values for a single similarity metric, 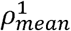 We repeated this procedure for the parcels identified as targets for sessions 2 and 3 to calculate 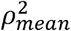 and 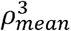, respectively. We then compared these RSFC profile similarity metrics to the “null” distribution of similarity metrics for all other parcels (N = 357) that were not associated with the TMS coil positions. We reasoned that 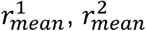, and 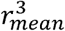 would differ from the null distribution if the TMS coil substantially influenced RSFC profiles for parcels centered beneath the coil.

We separately tested whether the coil impacted RSFC magnitude, because the RSFC profile similarity metric only describes the consistency of the *relative patterns* of RSFC. Conceivably, RSFC strength between parcel pairs could be impacted specifically for parcels near the TMS coil. To test this possibility, for each TMS target parcel (*T*), we computed the summed Euclidean distances between each session’s RSFC profile (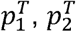, and 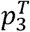) and the average RSFC profile over the sessions 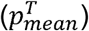 for a single error metric, 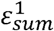, that summarizes the connectivity strength differences for each parcel over the sessions. We then calculated analogous error metrics for all other parcels not associated with the TMS coil positions for a “null” distribution of errors. We reasoned that 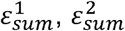, and 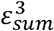would differ from the null distribution if the TMS coil substantially altered RSFC strengths for parcels centered beneath the coil.

## Results

### Static artifacts in phantom scans

Figure 2A shows the slice prescriptions for the PSF-EPI and conventional EPI phantom scans. Separate scans were conducted with the acquisitions arranged in parallel or oblique orientations with respect to the plane of the TMS coil. Visual inspection of slices acquired 15 mm (Fig. 2B; slice 5) and 30 mm (Fig. 2C; slice 10) from the TMS coil reveal spatial distortions and signal loss that qualitatively varied according to the EPI sequence, TMS coil presence, and acquisition orientation, with more substantial artifacts observed in the slice closer to the TMS coil. The raw images and SNR maps reveal slight distortions and signal loss in both EPI sequences even in the absence of the TMS coil. These patterns are exacerbated by introducing the TMS coil into the scanner. Notably, the PSF-EPI data contained considerably less distortion and less signal dropout relative to the conventional EPI data in both parallel and oblique acquisition orientations. We performed statistical analyses on image quality metrics calculated for the PSF-EPI and conventional EPI phantom scans to characterize the effects of the TMS coil and acquisition orientation quantitatively.

Structural images acquired using the 3D FLASH sequence are undistorted by the TMS coil. The asterisks indicate data acquired in the presence of the TMS coil. (C) Comparisons of the 10^th^ slice. Conventions as in *B*. R = right, L = left, S = superior, I = inferior, Phase = phase-encoding direction.

To compare TMS coil effects on signal strength, we calculated temporal SNR in each condition. Over all conditions, temporal SNR values were greater with PSF-EPI compared to conventional EPI in nearly all image slices (Fig. 3A). Elevated tSNR in the PSF-EPI was more evident when the TMS coil was present and in slices closer to the TMS coil (Fig. 3A). A rmANOVA conducted on the slice-averaged tSNR values (Fig. 3B) revealed significant main effects of TMS coil presence (F_1,36_ = 6.8, p = 0.01), sequence (F_1,36_ = 1218.7, p = 2.3e-29), and acquisition orientation (F_1,36_ = 48.7, p = 3.8e-8). These significant main effects reveal that the presence of the TMS coil reduced tSNR, the PSF-EPI sequence produced images with higher tSNR, and higher tSNR values were observed when the acquisitions were oriented parallel to the TMS coil. Significant interaction effects were observed between the TMS coil and sequence (F_1,36_ = 50.9, p = 2.2e-8), the TMS coil and acquisition orientation (F_1,36_ = 230.3, p = 3.2e-17), and the sequence and acquisition orientation (F_1,36_ = 90.9, p = 2.2e-11). Furthermore, the TMS coil X sequence X acquisition orientation interaction also achieved statistical significance (F_1,36_ = 74.3, p = 2.8e-10). Because of the significant interaction effects, we compared tSNR values achieved with PSF-EPI and conventional EPI in each condition (Fig. 3B) and found that tSNR values were significantly higher with PSF-EPI in all conditions (p < 2.2e-19 in all 4 paired t-tests).

**Fig. 3.**
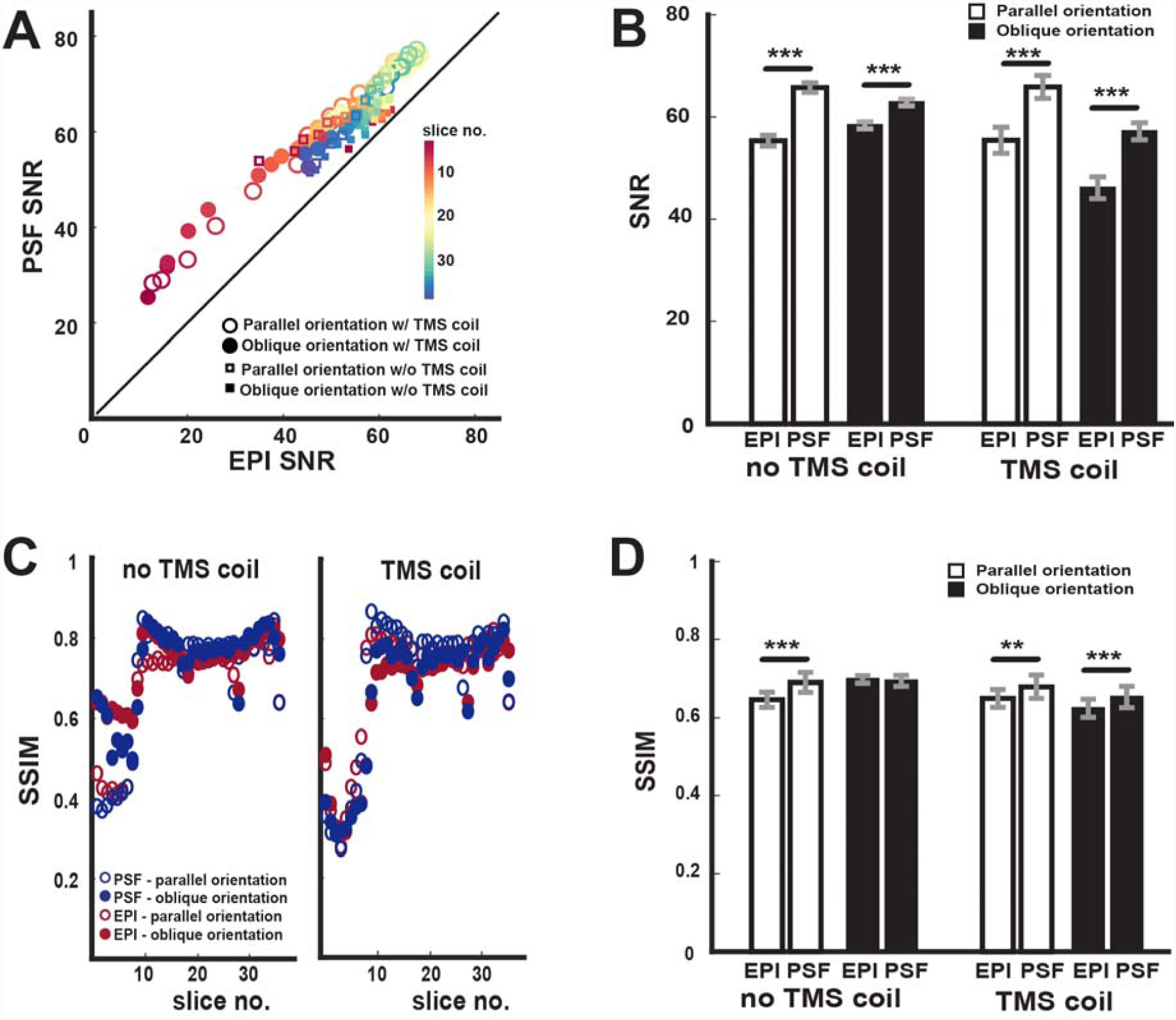
Quantitative comparisons between the conventional EPI and PSF-EPI data. (A) Markers indicate the average SNR values for each slice of the conventional EPI and PSF-EPI data acquired in the phantom scans calculated for parallel and oblique acquisitions, with and without the TMS coil. Points above the unity line indicate higher SNR with the PSF-EPI sequence. (B) Average SNR values with conventional EPI and PSF-EPI over the slices calculated for parallel and oblique acquisitions, with and without the TMS coil. (C) Markers indicate SSIM between the 3D FLASH images and the images acquired using conventional EPI and PSF-EPI. (D) Average SSIM values with conventional EPI and PSF-EPI over the slices calculated for parallel and oblique acquisitions, with and without the TMS coil. Error bars indicate SEM. ** p < 0.01; *** p < 0.001.

To compare TMS coil effects on image distortion, we calculated SSIM between the 3D FLASH phantom volume and the EPI volumes acquired under each condition. Larger SSIM values reflect greater similarity with the 3D FLASH data – these data are not susceptible to spatial distortions and signal loss induced by the TMS coil presence so they serve as a ground truth. Over all conditions (Fig. 3C), SSIM values were greater with PSF-EPI compared to conventional EPI in nearly all image slices. The presence of the TMS coil tended to reduce SSIM in slices close to the coil, particularly when images were acquired using an oblique orientation (Fig. 3C). A rmANOVA conducted on the slice-averaged SSIM values (Fig. 3D) revealed significant main effects of TMS coil presence (F_1,36_ = 9.2, p = 0.004) and sequence (F_1,36_ = 12.5, p = 0.001), but not of acquisition orientation (F_1,36_ = 0.01, p = 0.92). These significant main effects confirm that the presence of the TMS coil induces distortions in the EPI data and that the PSF-EPI sequence produced images more similar to the 3D FLASH images. Although the TMS coil X sequence interaction failed to achieve significance (F_1,36_ = 3.5, p = 0.07), significant 2-way interactions were observed between the TMS coil and acquisition orientation (F_1,36_ = 14.6, p = 0.0005) and the sequence and acquisition orientation (F_1,36_ = 23.4, p = 2.5e-5). Furthermore, the TMS coil X sequence X acquisition orientation interaction also achieved statistical significance (F_1,36_ = 28.3, p = 5.5e-6). Because of the significant 3-way interactions, we compared SSIM values achieved with PSF-EPI and conventional EPI in each condition (Fig. 3D). In the absence of the TMS coil, PSF-EPI yielded significantly higher SSIM values in the parallel acquisition orientation (p = 2.4e-5), but not with oblique acquisitions (p = 0.63). Importantly, SSIM values were significantly higher with PSF-EPI irrespective of orientation when the coil was present (paired t-test, parallel: p = 0.002; oblique: p = 0.0001).

### Static artifacts in brain scans

We compared how the placement of a TMS coil over left sensorimotor cortex impacted brain images acquired using PSF-EPI, conventional EPI, and SMS-EPI (Fig. 4A). Image distortions and ghosting artifacts were obvious in visual inspections of slices near the TMS coil, particularly in the conventional EPI and SMS-EPI data (Fig. 4B). Whole-brain tSNR maps (Fig. 5) clearly reveal substantial signal loss in brain regions under and near the TMS coil in the conventional EPI and SMS-EPI scans. In contrast, the tSNR map computed for the PSF-EPI scan appears far more homogeneous.

**Fig. 4.**
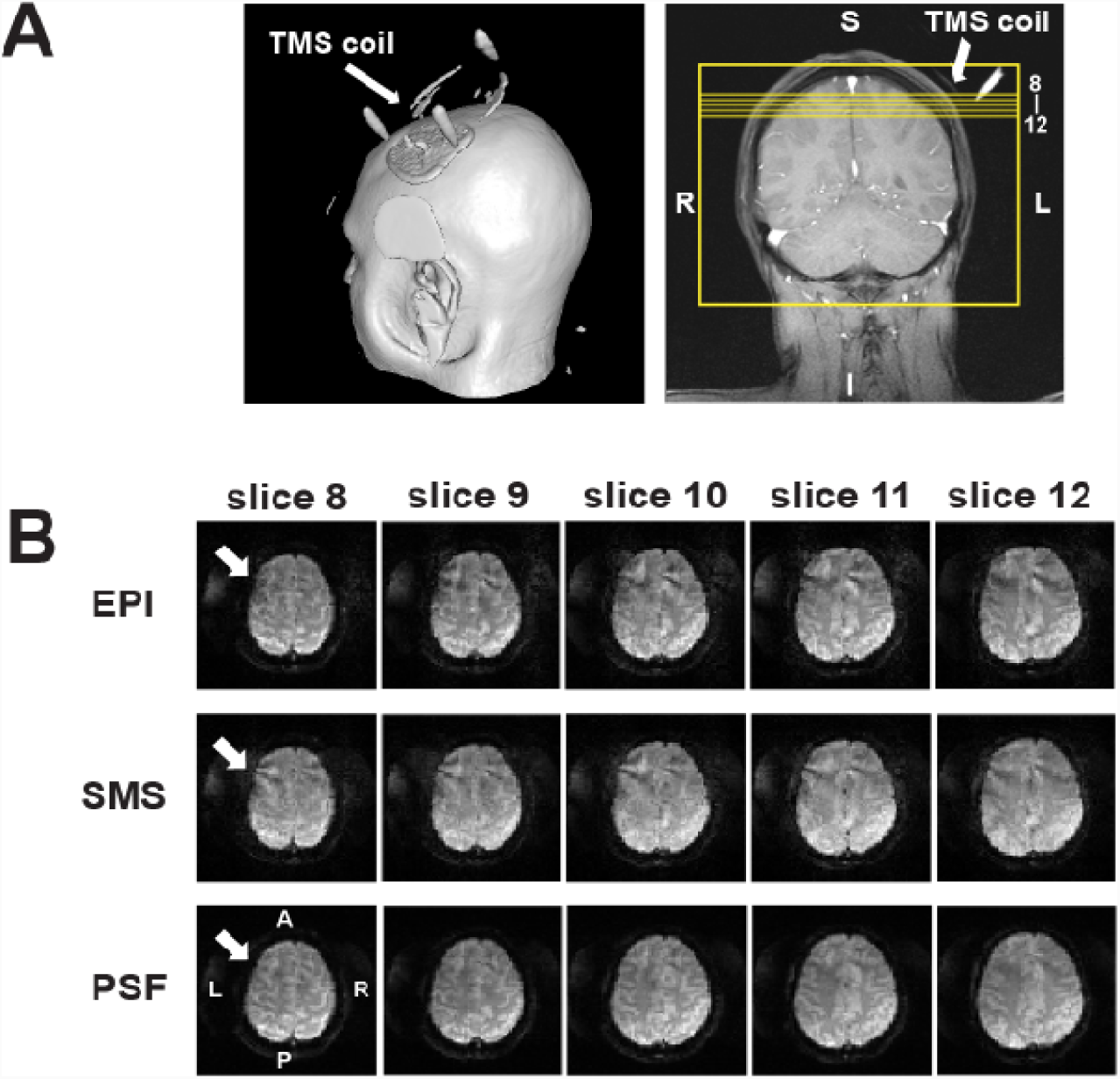
Comparison of raw images acquired in the presence of the TMS coil. (A) Left: Volume rendering depicting the position of the TMS coil with respect to the scalp. Right: Slice prescriptions used for PSF-EPI, conventional EPI, and SMS scans. (B) Images acquired with the conventional EPI, SMS, and PSF-EPI sequences in the presence of the TMS coil (white arrow). Distortion and ghosting artifacts are visible in slices in close proximity to the TMS coil. L = left, R = right, A = anterior, P = posterior.

**Fig. 5.**
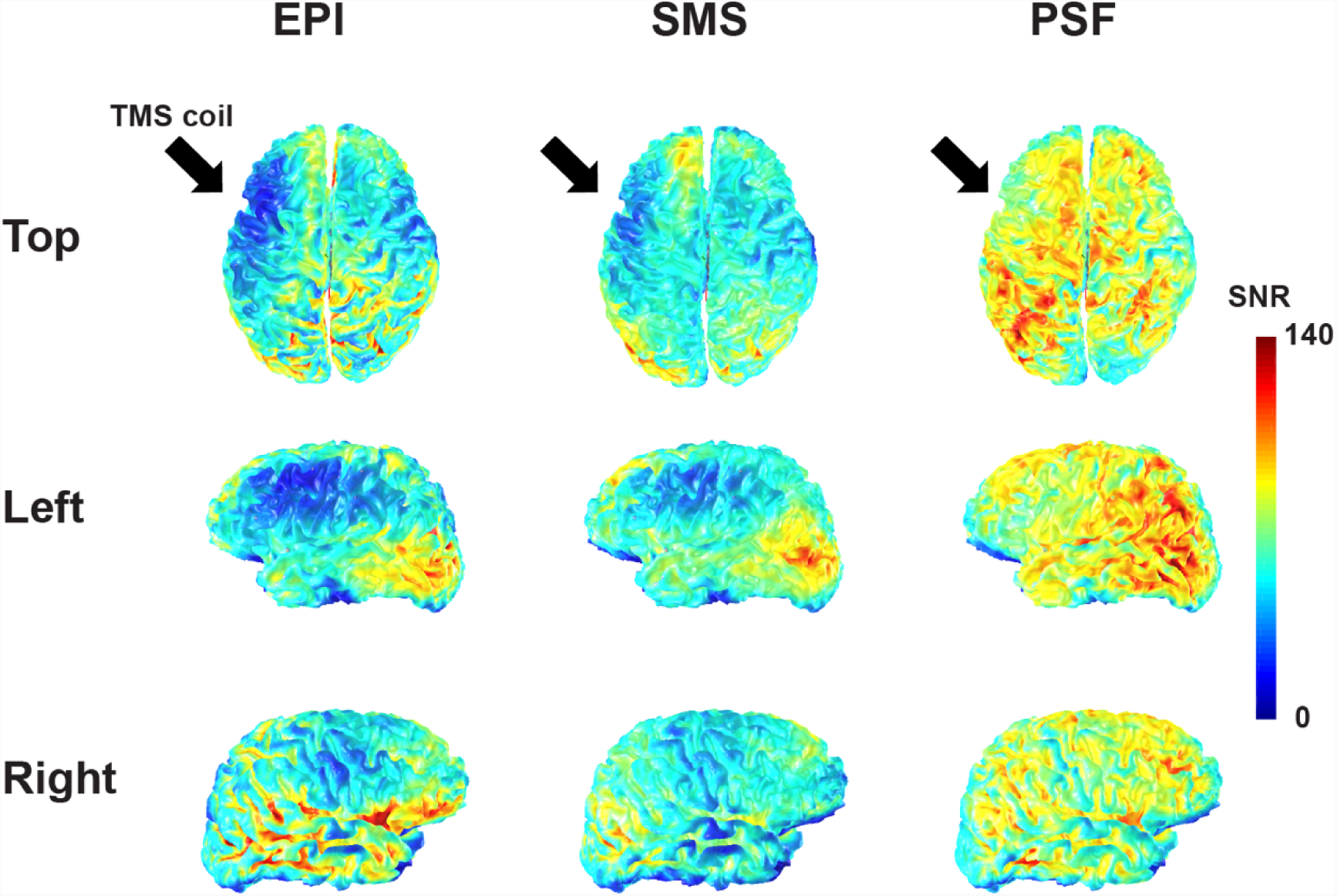
SNR maps computed for conventional EPI, SMS, and PSF-EPI. The black arrow indicates the TMS coil location (see Fig. 4). The TMS coil introduces an obvious shadow in the conventional EPI and SMS data, which is less obvious with PSF-EPI.

Quantitative comparisons between the EPI sequences are shown in Figure 6. Mean tSNR values of the PSF-EPI data exceeded those of the conventional EPI and SMS-EPI data in most slices (Fig. 6A). A rmANOVA conducted on the slice-averaged tSNR values (Fig. 6B) revealed a significant main effect of sequence (F_2,72_ = 248.0, p = 5.1e-33). Post-hoc tests revealed that the PSF-EPI sequence yielded higher tSNR compared to conventional EPI (p = 3.6e-10) and SMS-EPI (p = 1.3e-19). Conventional EPI also yielded higher tSNR compared to SMS-EPI (p = 5.0e-18). These results indicate that PSF-EPI achieves greater tSNR in brain data acquired in the presence of the TMS coil compared to conventional EPI and SMS-EPI.

**Fig. 6.**
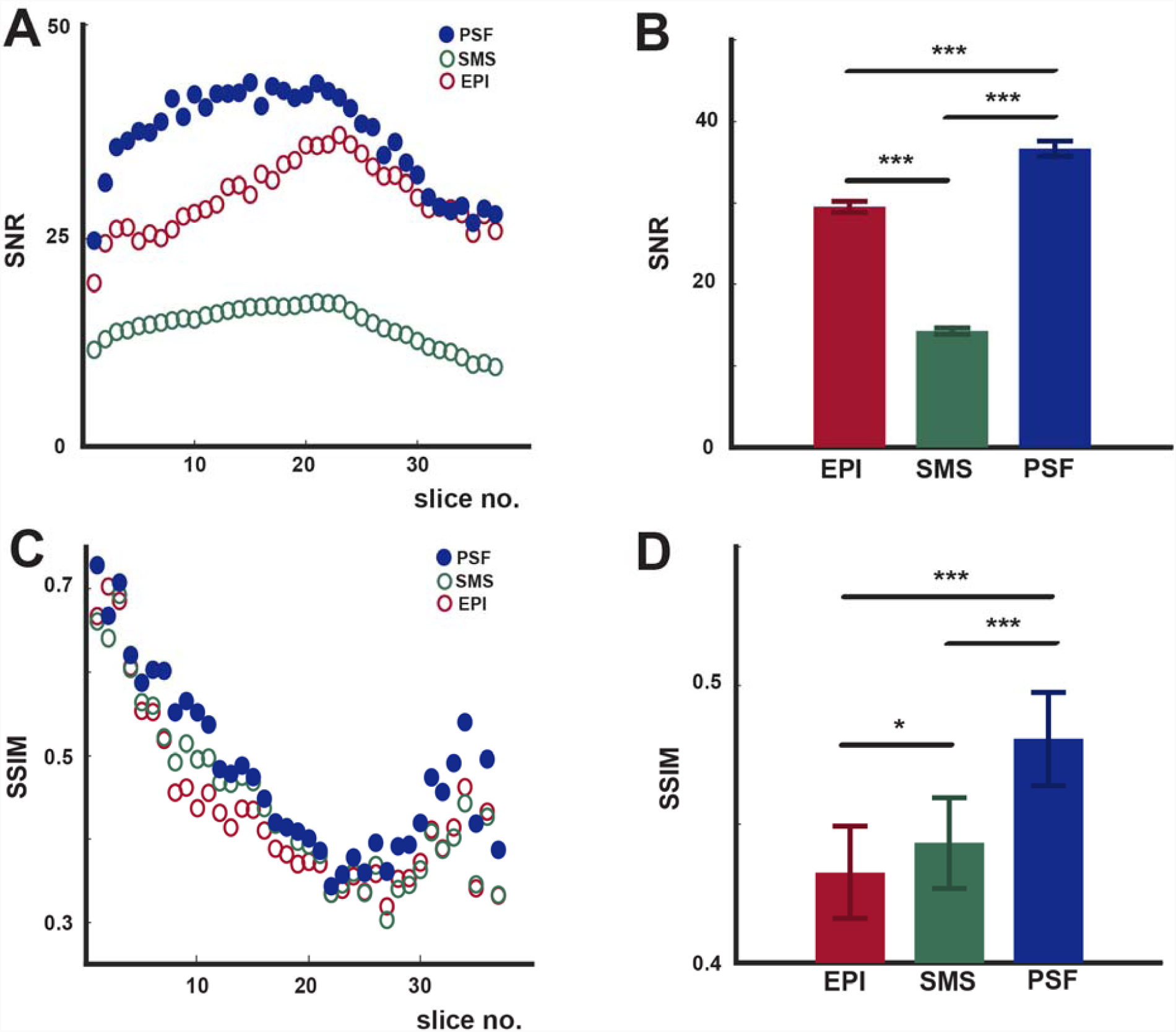
Quantitative comparisons between the conventional EPI, SMS and PSF-EPI data. (A) Markers indicate the average SNR values for each slice acquired with the PSF-EPI, conventional EPI, and SMS sequences. (B) Comparison between the average SNR values of the conventional EPI, SMS, and PSF-EPI data averaged over slices. (C) SSIM between the 3D FLASH images and the images acquired using conventional EPI, SMS, and PSF-EPI. (D) Comparison between the SSIM values achieved with the conventional EPI, SMS, and PSF-EPI sequences averaged over all slices. Error bars indicate SEM. * p < 0.05; *** p < 0.001.

Quantitative comparisons between the SSIM computed for each EPI sequence similarly indicated the superiority of PSF-EPI. Across slices, SSIM tended to be higher for PSF-EPI compared to conventional EPI and SMS-EPI (Fig. 6C). A rmANOVA conducted on the slice-averaged SSIM values (Fig. 6D) revealed a significant main effect of sequence (F_2,72_ = 63.6, p = 1.2e-16). Post-hoc tests revealed that SSIM was higher with PSF-EPI compared to conventional EPI (p = 8.5e-12) and SMS-EPI (p = 8.0e-10). SSIM was higher with SMS-EPI compared to conventional EPI (p = 0.01).

### Acute artifacts in phantom scans

To characterize the temporal profile of TMS pulse effects on image acquisition using PSF-EPI, we delivered a single TMS pulse at different times preceding and during volume acquisition (Fig. 7A). To quantify TMS effects as a function of delay times (i.e., the interval between the pulse and image acquisition), we calculated the root-mean-square error (RMSE) between a volume acquired before TMS delivery and volumes acquired during or immediately following TMS on a slice-wise basis (Materials and methods). Because interactions between the scanner and TMS coil can depend on the orientation of the coil with respect to the B_0_ field (Yau et al., 2014), we characterized acute TMS effects on PSF-EPI data with the coil aligned with or perpendicular to the B_0_ field (Fig. 7B). RMSE profiles with TMS intensity set at 70% and 100% maximum stimulator output. For both coil positions, signal perturbations indexed by RMSE were observed mostly in slices close to the TMS coil which were acquired earlier in the slice sequence (Fig. 7C). In general, TMS applied during acquisition (0ms and 50ms with respect to acquisition onset) induced the largest image perturbations. When the TMS coil was aligned to the B_0_ field, TMS preceding acquisition also perturbed images and errors were observed when image acquisition followed TMS pulse delivery by less than 75 ms (Fig. 7C). In this coil position, increasing TMS intensity from 70% to 100% maximum output resulted in larger errors at the longer delays and disrupted signal quality in more slices (Fig. 7C,D). When the TMS coil was oriented perpendicular to the B_0_ field, TMS pulses preceding image acquisition had little effect on RMSE and comparable distortions were observed with the 70% and 100% TMS intensities. The fact that different spatiotemporal RMSE profiles were observed for the different TMS coil orientations (Fig. 7C,D) confirms that interactions between the scanner and TMS vary according to the relative orientation of the coil with respect to the B_0_ field. These differences may also reflect, in part, the slice prescriptions used under the two coil orientations: Subtle differences in SNR could conceivably result in apparent differences in vulnerability to the eddy current produced by TMS. Critically, volumes acquired >3 s after TMS delivery did not contain substantial errors even when the TMS pulse potentially coincided with RF excitation and image acquisition (Fig. 7D).

**Fig. 7.**
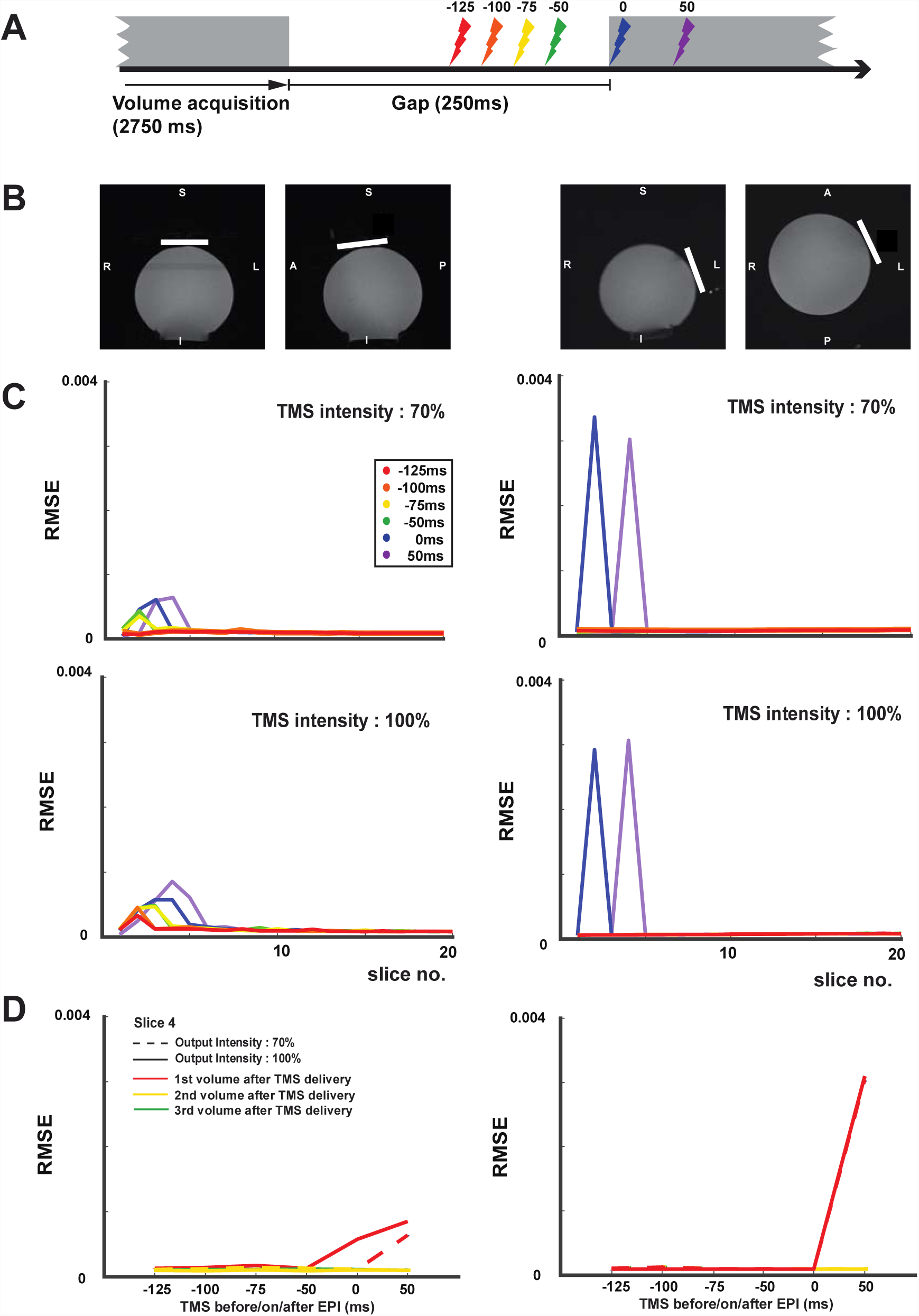
Characterization of acute TMS artifacts. (A) Diagram indicates potential times a single TMS pulse was delivered with respect to image acquisition (Materials and methods). (B) The TMS coil (white bar) was placed superior to the phantom with its plane was aligned to the transverse imaging plane (Two left images in coronal and sagittal view) and approximately left lateral to the phantom (Two right images in coronal and axial view). In these configurations, the field generated by the coil was either aligned with or perpendicular to the B_0_ field, respectively. (C) The root mean square error (RMSE) values calculated for each slice indicate the differences between reference images acquired prior to TMS and images acquired with TMS delivered at different times with respect to image acquisition. For each coil position (superior = left column; lateral = right column), separate RMSE profiles are indicated for TMS intensity set at 70% (upper panels) and 100% (lower panels) maximum stimulator output. (D) RMSE for the 4^th^ slice (∼12 mm from the TMS coil) with the stimulator output 70% (dashed line) and 100% (solid line) as a function of TMS delivery time. Red, yellow, and green traces indicate 1^st^, 2^nd^, and 3^rd^ volume during or after TMS delivery, respectively. RMSE temporal profiles differed slightly depending on whether the TMS was positioned superior (left) or lateral (right) to the phantom. In both positions, RMSE increased in volumes acquired during or immediately after TMS delivery (red), but returned to baseline levels in volumes separated from the TMS pulse by intervals exceeding 3 s (yellow and green). Error bars indicate SEM.

### Simultaneous TMS-fMRI experiment

In one participant, we characterized BOLD signal changes associated with TMS and performance of a finger tapping task to validate the use of PSF-EPI (Fig. 8A). The goal of this experiment was simply to verify that BOLD signal changes could be measured in brain regions in the vicinity of the TMS coil and regions removed from the TMS coil. In separate block design scans, bursts of low-or high-intensity TMS were delivered between volume acquisitions during each block. Subjects also performed bimanual finger tapping, cued by low-intensity TMS, during one scan. The TMS coil was positioned over left sensorimotor cortex (Materials and methods). Bimanual finger tapping paired with TMS_Low_ was associated with robust BOLD signal changes in a distributed sensorimotor network including the bilateral precentral gyrus, bilateral postcentral gyrus, bilateral cerebellum, left inferior parietal and right middle temporal gyrus (Fig. 8B). In order to compare BOLD signals in corresponding sensorimotor regions near to and removed from the TMS coil, we identified the peak activations associated with bimanual finger tapping in the left (x = −40, y = −19, z = 56) and right precentral gyri (x = 44, y = −22, z = 59) and defined two spherical regions-of-interest (ROI; 5-mm radius) for extracting response time series and activation estimates (Fig. 8B). The response equivalence between the two sensorimotor ROIs indicates that the PSF-EPI sequence successfully preserved BOLD signal changes associated with finger tapping in brain regions in the vicinity of the TMS coil. TMS alone was also associated with BOLD signal changes in the distributed sensorimotor network identified with finger tapping (Fig. 8C), with qualitatively greater BOLD signal changes in the high-intensity TMS condition compared to the low-intensity condition. Although BOLD signal changes related to TMS were weaker than those associated with finger tapping, the three conditions were marked by highly similar temporal response profiles.

**Fig. 8.**
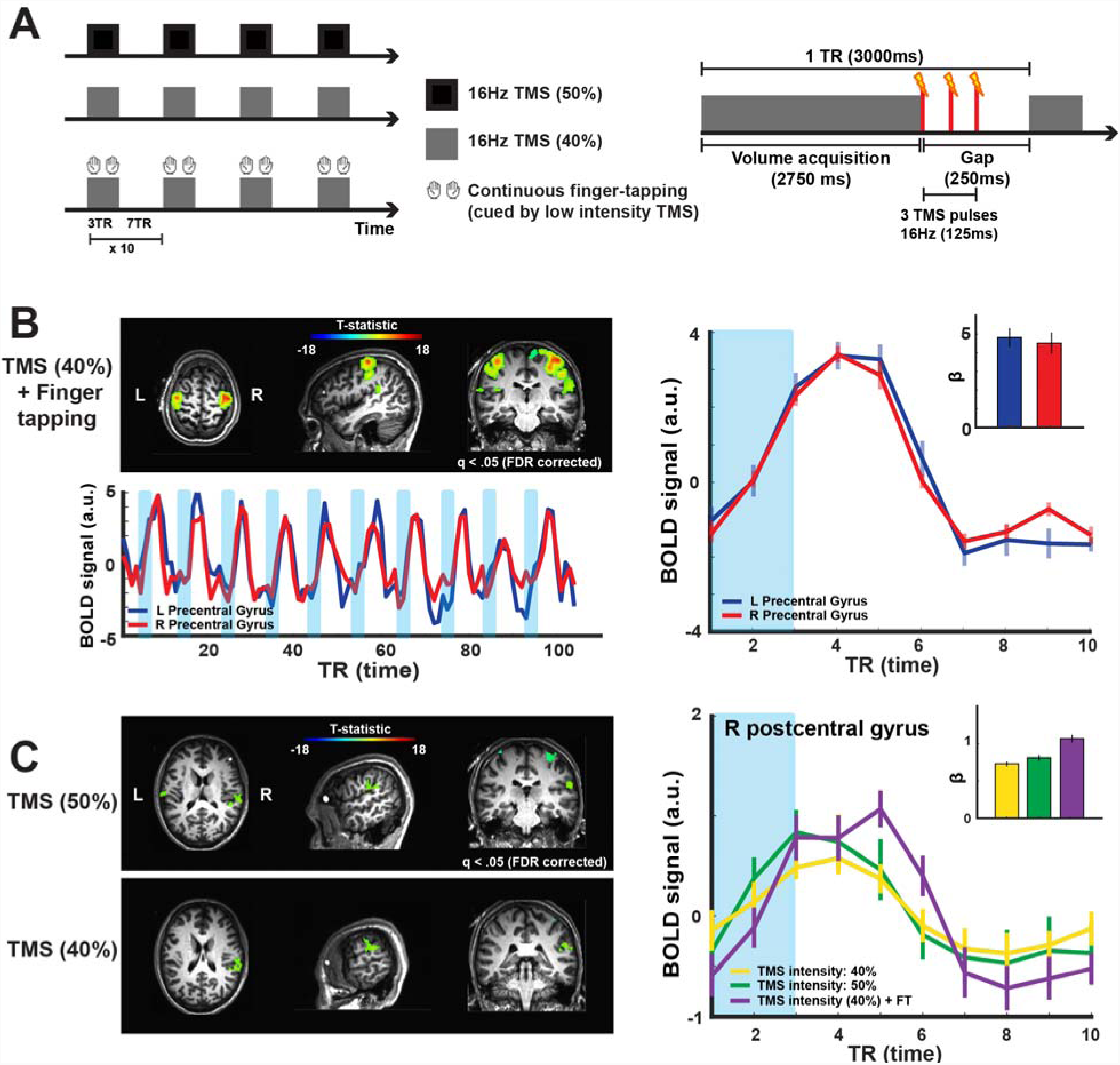
Simultaneous TMS-fMRI scans. (A) In separate block-design scans, a subject received low-and high-intensity TMS bursts (40% and 50% maximum stimulator output, respectively; Materials and methods). In a separate scan, the subject also performed bimanual finger tapping when cued by low intensity TMS. TMS bursts (triplet; 16Hz) were applied during a 250-ms gap separating consecutive volume acquisitions. (B) Activation pattern associated with bimanual finger tapping and low intensity TMS. BOLD signal time series from the left and right precentral gyri reveal robust signal changes associated with finger tapping. The temporal response profiles and response magnitudes in the left and right hemispheres were comparable, indicating that the TMS coil (positioned over the left hemisphere) did not substantially attenuate signal locally. (C) Activation patterns associated with low-and high-intensity TMS alone. The temporal response profiles were highly similar for the finger-tapping and TMS conditions. Error bars indicate SEM

### Resting state functional connectivity

To test whether the PSF-EPI sequence could be used to acquire resting state fMRI data in the presence of the TMS coil, we conducted resting state fMRI scans with one subject over 3 sessions. Because the TMS coil was positioned over different scalp locations in each session (Materials and methods), we could characterize the consistency of RSFC patterns across the sessions and test whether the positioning of the TMS coil (Fig. 9A) specifically affected resting state networks comprising regions underneath the coil. We found that RSFC patterns over the whole brain (Fig. 9B), sorted according to an existing parcellation scheme (Glasser et al., 2016), were highly similar over sessions (mean r = 0.78; range: 0.75-0.80). These correlations are comparable to published estimates of across-session RSFC consistency comparing 15-min portions of resting state data acquired with conventional EPI at 3T (Laumann et al., 2015; Xu et al., 2016; Gordon et al., 2017). We evaluated the across-session similarity of RSFC patterns for the parcels identified as the TMS targets (Materials and methods) and found that these values (target 1: 0.86, target 2: 0.84, target 3: 0.81) fell well within the distribution of average similarity values for all parcels not associated with TMS coil position (Fig. 9C; mean of null distribution: 0.77). Similarly, we evaluated the summed differences in RSFC strength (i.e., Euclidean distances, see Materials and methods) for the TMS target regions (target 1: 0.18, target 2: 0.18, target 3: 0.20) and found that these values also fell well within the null distribution defined by parcels not associated with TMS coil position. Together these, results indicate that the PSF-EPI sequence can produce high quality RSFC data in the presence of the TMS coil.

**Fig. 9.**
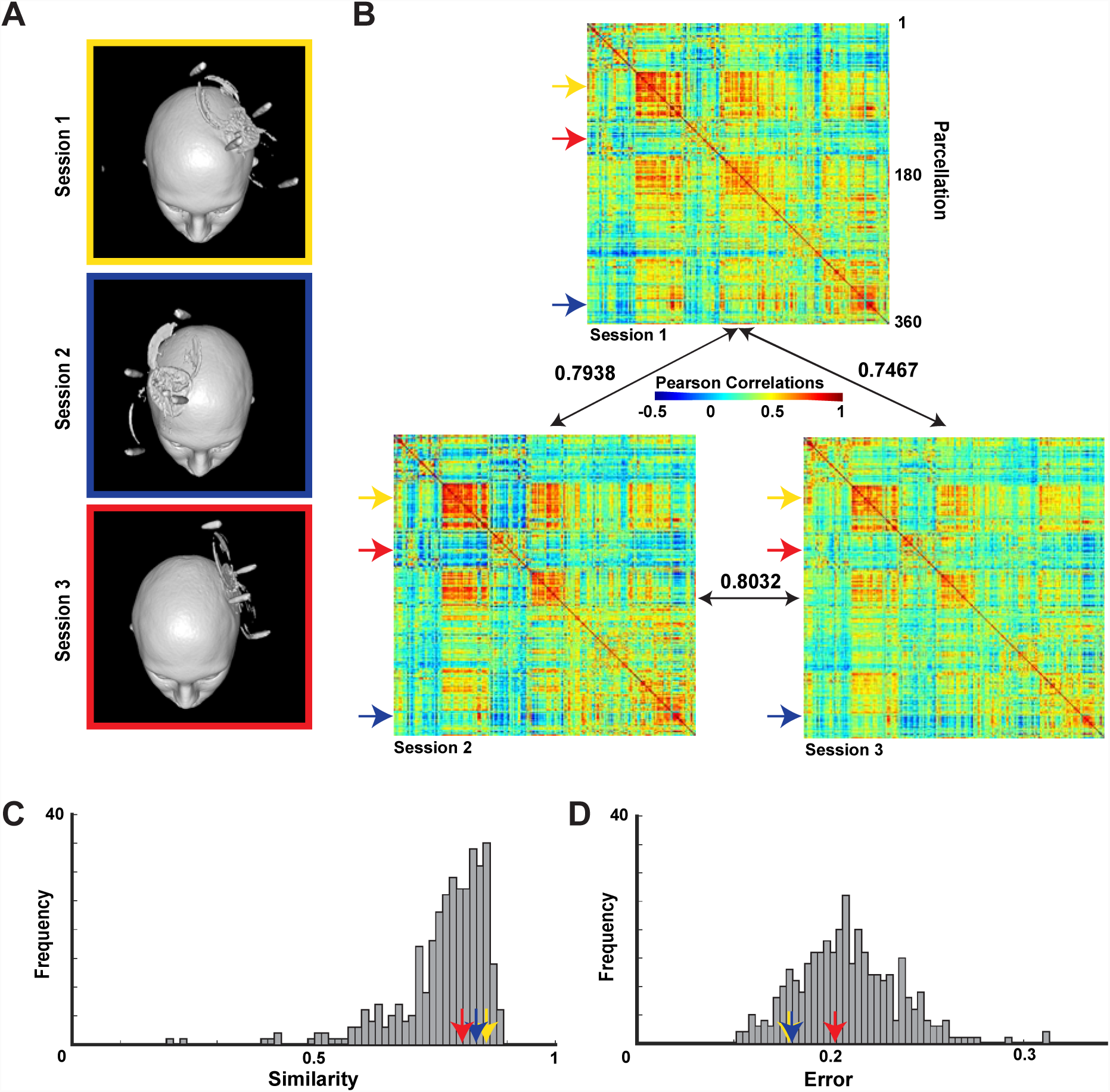
Resting state functional connectivity measured in the presence of the TMS coil. (A) Resting state fMRI data was acquired from a single subject in 3 sessions (15 minutes of data per session). The TMS coil was positioned over different scalp locations in each session. (B)Resting state functional connectivity (RSFC) matrices (Materials and methods) are depicted for 360 parcels spanning both hemispheres. The similarity of the RSFC patterns are indicated by the black arrows and associated correlations. The yellow, blue, and red arrows indicate the parcels associated with the TMS coil locations in sessions 1, 2, and 3, respectively. (C) The arrows indicate the average similarity in the RSFC patterns for the TMS-associated parcels (see B for reference). The grey bars indicate the null distribution of RSFC similarity values for all parcels not associated with the TMS coil positions. (D) The arrows indicate the total differences in the RSFC magnitudes for the TMS-associated parcels (see *B* for reference). The grey bars indicate the null distribution for all parcels not associated with the TMS coil positions.

## Discussion

We sought to demonstrate the feasibility of using PSF-EPI to reduce metal-induced susceptibility artifacts and signal dropouts in simultaneous TMS-fMRI experiments. In phantom and brain scans, we found that the PSF-EPI sequence yielded data that were marked by less spatial distortion, ghosting artifacts, and signal dropout compared to data acquired using conventional EPI and SMS-EPI. We then characterized the temporal profile of acute artifacts related to TMS discharge and established that 100ms were required between the TMS pulse and subsequent image acquisition to avoid substantial signal loss and image perturbations. We combined TMS with performance of a bimanual motor task to confirm that robust BOLD signal changes could be measured using PSF-EPI in brain regions near and removed from the TMS coil. Lastly, we demonstrated that high quality resting state fMRI data could be acquired using the PSF-EPI sequence even in the presence of the TMS coil.

Our data confirm that the introduction of a TMS coil to the scanner environment can result in substantial image artifacts and signal loss in EPI data. Note that we only tested a single TMS system and a particular combination of receiver coils, so our results may not generalize to setups comprising other TMS equipment or coil arrangements. That said, in our setup, we found in phantom scans that, while the PSF-EPI sequence did not entirely overcome the artifacts, spatial distortions and signal loss were significantly reduced with PSF-EPI compared to conventional EPI in each of our tested conditions. PSF-EPI also yielded data with less distortions and signal loss in brain scans compared to conventional EPI and SMS-EPI. Moreover, consistent with previous reports (Bestmann et al., 2003a), we found that signal loss and distortions with conventional EPI depended on the orientation of the slice prescriptions with respect to the plane of the TMS coil: Greater artifacts are present when the frequency encoding direction is oblique rather than parallel to the plane of the TMS coil. Accordingly, previous studies using conventional EPI have attempted to reduce the coil-related static artifacts by prescribing the image acquisitions with respect to the TMS coil arrangement (Bestmann et al., 2003a; Moisa et al., 2009; Weiskopf et al., 2009; Bungert et al., 2012; Navarro de Lara et al., 2015, 2017). Although we found that the PSF-EPI sequence, like the conventional EPI sequence, was sensitive to the slice acquisition orientation with respect to the TMS coil in phantom scans, the PSF-EPI sequence yielded higher quality data in all orientations according to tSNR and SSIM estimates. Consistent with these phantom scan results, we observed that PSF-EPI sequence yielded substantially better signal and less distortion compared to conventional EPI and SMS sequences even as we used transverse prescriptions in our brain scans. We also used the transverse prescription in the simultaneous TMS-fMRI and resting state fMRI scans and observed robust BOLD signal changes and consistent RSFC patterns despite the fact that we did not tailor the image prescriptions to the TMS coil orientation and placement. Thus, in simultaneous TMS-fMRI experiments where coil-related artifacts are present and concerning, the use of PSF-EPI is advantageous over conventional EPI based on image quality metrics as well as ease and flexibility of use.

In the phantom and brain scans, we also observed substantial ghost artifacts along the phase encoding direction with conventional EPI which are presumably caused by systematic inconsistencies between odd and even k-space lines in the acquired data (Hu & Le., 1996). These ghost artifacts have also been reported in previous simultaneous TMS-fMRI experiments (Bestmann et al., 2003a; Bungert et al., 2012). Earlier studies using conventional EPI dealt with this artifact by oversampling in the phase encoding direction, which shifted the ghost artifacts caused by the TMS coil (Blankenburg et al., 2008; Ruff et al., 2008; Feredoes et al., 2011). This strategy, though, can result in a subsequent decrease in SNR and an increase in phase errors (Hu & Glover., 2006). Using PSF-EPI, we observed qualitatively less ghosting, consistent with earlier studies that reported reductions in ghost artifacts through the use of PSF-EPI sequences (Zaitsev et al., 2004; Chung et al., 2011).

We characterized the acute artifacts associated with discharging a single TMS pulse in order to identify the optimal delay between TMS and subsequent image acquisition. Depending on the hardware setup, the introduction of this delay can be important for allowing the eddy currents associated with TMS discharge to clear without impacting subsequent RF excitation and image acquisition. Because interactions between TMS and the scanner can depend on the relative orientation of the TMS coil with respect to the B_0_ field, we characterized the acute artifacts associated with a TMS pulse with the coil oriented parallel or perpendicular to the B_0_ field. When the TMS coil was aligned with the B_0_ field, we found that image quality remained high with delay times exceeding 50 ms. This timing is consistent with a recent report that delays on the order of 50 ms are sufficient to avoid distortions in the images with the 100% TMS output intensity (Navarro de Lara et al., 2017). Notably, we observed a different spatiotemporal perturbation pattern when the TMS coil was perpendicular to the B_0_ field (Fig. 7). In this orientation, pulse-related distortions were only observed when image acquisition occurred during or immediately following the TMS pulse (i.e., TMS pulse time at 0 and +50 ms relative to image acquisition). The different disruption patterns we observed confirm the sensitivity of TMS effects in the scanner environment to the relative orientation of the static field and highlight the variability in acute TMS influences on image quality even when using the same hardware. Importantly, while we found that image quality was minimally affected by TMS at delays exceeding 50 ms, some studies have recommended using longer delays on the order of 100 ms (Shastri et al., 1999; Bestmann et al., 2003a). This variation across studies is likely due to the use of different TMS stimulators and coils, different scanners and imaging hardware, and different EPI sequences. Indeed, TMS-related eddy currents may minimally affect image quality with some stimulator and coil designs. Moreover, some acute temporal artifacts related to TMS conceivably may be a consequence of imprecise coordination between the scanner and TMS system so better synchronization (e.g., timing control via a dedicated microprocessor) may also reduce TMS-related artifacts. Because acute TMS artifacts may differ dramatically across setups, researchers should carefully determine whether acute artifacts are present in their experiment and introduce a delay between TMS pulsing and image acquisition as needed.

We validated the feasibility of the PSF-EPI sequence in simultaneous TMS-fMRI experiments by characterizing TMS and motor responses. Our goal in this experiment was simply to confirm that systematic BOLD signal changes could be measured over the brain rather than to characterize activation patterns and infer network organization. Using PSF-EPI, we observed robust BOLD signal changes associated with bimanual finger-tapping in bilateral regions comprising a well-established distributed sensorimotor network. Importantly, we confirmed the location of the TMS over coil left motor cortex using our marker processing procedure and found that finger-tapping BOLD responses in sensorimotor regions beneath and near the coil were just as robust as responses in homologous regions in the hemisphere contralateral to the coil. This response equivalence is important for indicating that the BOLD signal in the vicinity of the coil was largely preserved with PSF-EPI, particularly because there has been some debate regarding the ability to measure significant BOLD signal changes in brain regions beneath the TMS coil. Indeed, many studies have failed to measure significant BOLD signal changes in motor cortex when delivering subthreshold TMS that successfully evokes significant activations in regions remote from the coil (Bohning et al., 1998; Baudewig et al., 2001; Kemna and Gembris, 2003; Bestmann et al., 2004). We also observed BOLD signal changes associated with TMS even in the absence of motor performance in distributed brain regions. These activation patterns, found in sensorimotor cortex, parietal operculum, and auditory regions, are largely consistent with the results of previous TMS-fMRI studies targeting on motor cortex (Bestmann et al., 2004; Moisa et al., 2009; Shitara et al., 2011; Navarro de Lara et al., 2017; Yau et al., 2013). Importantly, the responses to TMS were characterized by robust and systematic temporal response profiles that closely resemble the response time courses related to motor performance. Although the results from our feasibility experiment are insufficient for addressing specific questions related to network interrogation, the results do allow us to confirm that PSF-EPI can be used to characterize BOLD signal changes in simultaneous TMS-fMRI scans in distributed regions near to and removed from the TMS coil.

We demonstrated the feasibility of using PSF-EPI for acquiring resting state fMRI data in the presence of TMS coil. We confirmed that RSFC patterns were robust and highly correlated across sessions when comparing 15 min of data per session, consistent with previous reports (Laumann et al., 2015; Xu et al., 2016; Gordon et al., 2017). We additionally confirmed that RSFC patterns for brain regions under the TMS coil were reliable and robust. These results highlight the potential for combining simultaneous TMS-fMRI and resting state fMRI in the same session; the combined use of these complementary approaches may reveal novel insights regarding the intrinsic architecture of the human brain (Fox et al., 2014; Hawco et al., 2018) and understanding how remote TMS effects relate to RSFC may inform clinical interventions utilizing targeted brain stimulation (Fox et al., 2012, 2013).

In summary, we have characterized the static artifacts induced by the introduction of a TMS coil into the scanner environment. Relative to T1-weighted scans, scans acquired using PSF-EPI, conventional EPI, and SMS-EPI all showed some degree of image distortion and signal loss. Critically, data acquired using PSF-EPI suffered significantly less from the mere presence of our TMS coil, indicating that the characterization of the EPI distortions using point spread function mapping (Zaitsev et al., 2004; In and Speck, 2012; In et al., 2015, 2017b) and the subsequent adjustment of the EPI sufficiently corrected for the substantial magnetic field inhomogeneities induced by our TMS coil. Because PSF-EPI corrects for image distortion in a manner that is relatively invariant to changes in the prescription of the acquired volumes, its use enables flexible TMS coil positioning without the need to tailor prescriptions according to the specific coil placement and orientation. We additionally found that PSF-EPI yields functional data with minimal acute artifacts due to TMS discharge when the TMS pulse precedes RF excitation and image acquisition with a sufficient temporal gap. Based on the characterization of the static and acute artifacts, we conducted a simultaneous TMS-fMRI scan and found robust BOLD signal changes in distributed brain regions. Our collective results indicate that simultaneous TMS-fMRI experiments could benefit from the correction of EPI distortions using point spread function mapping. Importantly, some MRI-compatible TMS systems may not induce comparable spatial distortions and signal loss as we observed, so our current results may not generalize to all simultaneous TMS-fMRI setups. Nevertheless, in the event that the TMS system does introduce problematic spatial artifacts and signal loss, PSF-EPI may be helpful particularly when specialized hardware solutions for treating susceptibility effects are unavailable. PSF-EPI may also be a general solution for multimodal stimulation and imaging studies (e.g., EEG-fMRI (Liu et al., 2006; Abreu et al., 2018), tDCS-fMRI (Callan et al., 2016), and focused ultrasound-fMRI (Legon et al., 2014) that require the introduction of metallic hardware into the scanner environment. Moreover, PSF-EPI may hold great promise for measuring functional connectivity from the combination of simultaneous TMS-fMRI and resting state fMRI (Fox et al., 2012).

## Acknowledgements

This research was supported by R01NS097462 and The Dana Foundation. We thank Gisela Hagberg for providing the PSF-EPI sequence. We thank Meghan Robinson for assisting with the resting state analysis and field map characterization. We thank Yau Lab members for thoughtful discussions. This work was performed in BCM’s Core for Advanced MRI (CAMRI).

